# Scalable sequence database search using Partitioned Aggregated Bloom Comb-Trees

**DOI:** 10.1101/2022.02.11.480089

**Authors:** Camille Marchet, Antoine Limasset

## Abstract

The Sequence Read Archive public database has reached 45 Peta-bytes of raw sequences and doubles its nucleotide content every two years. Although BLAST-like methods can routinely search for a sequence in a small collection of genomes, making searchable immense public resources accessible is beyond the reach of alignment-based strategies. In recent years, abundant literature tackled the task of finding a sequence in extensive sequence collections using *k*-mer-based strategies. At present, the most scalable methods are approximate membership query data structures that combine the ability to query small signatures or variants while being scalable to collections up to 10,000 eukaryotic samples. Here, we present PAC, a novel approximate membership query data structure for querying collections of sequence datasets. PAC index construction works in a streaming fashion without any disk footprint besides the index itself. It shows a 3 to 6 fold improvement in construction time compared to other compressed methods for comparable index size. A PAC query can need single random access and be performed in constant time in favorable instances. Using limited computation resources, we built PAC for very large collections. They include 32,000 human RNA-seq samples in five days, the entire Genbank bacterial genome collection in a single day for an index size of 3.5TB. The latter is, to our knowledge, the largest sequence collection ever indexed using an approximate membership query structure. We also showed that PAC’s ability to query 500,000 transcript sequences in less than an hour. PAC’s open-source software is available at https://github.com/Malfoy/PAC.

## 1 Introduction

Public databases such as the SRA (Sequence Read Archive) or ENA (European Nucleotide Archive) overflow with sequencing data. The vast amount of sequences, experiments, and species allows, in principle, ubiquitous applications for biologists and clinicians. Such databases are becoming a fundamental shared resource for daily sequence analysis. But concretely, we only witness the onset of their exploitation, while their exponential growth poses serious scalability challenges (for instance, SRA has reached 45 petabytes of raw reads and roughly doubles every three years [1]).

Today, searching for a query sequence in a single genome / dataset or a restricted collection of genomes is considered routine through alignment-based tools such as BLAST [4] and others [21, 19, 15, 11]. In contrast, the scale of databases such as SRA makes BLAST and any alignment-based method ill-suited due to the prohibitive cost of the alignment phase. Broader usages of such resources necessitate algorithms allowing hyper-scalable membership queries as a foundation. In particular, they must be able to quickly discard a sequence that is absent from a substantial dataset collection. They must also identify the datasets where the query is present. Therefore, recent literature has tackled these problems using alignment-free *k* -mer-based strategies.

The known most scalable solutions are sketching methods, which reduce the datasets to a small set of signatures based on the principle of locality-sensitive hashing. However, the loss of resolution implied by these methods narrows the query possibilities to large queries of the order of magnitude of genomes or larger. Numerous applications rely on significantly smaller query size, e.g. variant calling of SNP/small indels, finding alternative splicing sites, or other small genomic signatures.

Recent alignment-free literature has described novel methodologies to fill a dual need: small queries in vast dataset collections. Mostly, two paradigms [23] cover this question, both considering each dataset and the query itself as *k*-mer sets. The first type of approach relies on exact *k*-mer set representations. These methods build an associative index (using hash tables [3, 17, 24, 28] or FM-indexes [12, 27, 5]) where the key set corresponds to all *k*-mers of the collection. SUch static structure hardly scale to extensive instances with large *k*-mer cardinality but can be used for queries with high precision or to build de Bruijn graphs. Some indexing implement some kind of dynamicity either via merging or using dynamic bit vectors [18, 26, 2].

The second family of methods relies on probabilistic set representations [31, 16, 9, 6]. These approximate membership query (AMQ) structures trade false positives during the query for improved speed and memory performance compared to the previous category.

Approximate membership query approaches allowed the indexing of hundreds of thousands of microbial datasets [7]. For mammalian datasets, scaling to more samples is a timely challenge since the increase in the content of *k*-mer / datasets poses serious issues for the construction/footprint of current methods. Thus, to our knowledge, no published method has yet overcome the barrier set by SeqOthello [33] of indexing over 10,000 mammalian RNA samples.

This manuscript proposes a novel approximate membership query method to query biological sequences in collections of datasets dubbed PAC (Partitioned Aggregative Bloom Comb Trees). In our application case, queries are typically alternative splicing events or a short genomic context around a small variant or mutation, for both eukaryotic and microbial species. Our structure is designed to be highly scalable with the number of indexed samples and requires moderate resources.

## 2 Methods

In this section, we present our Partitioned Aggregative Bloom Comb Trees. We kept PAC as an acronym that covers the three keywords, including the main novelties compared to previous tree structures. PAC is available at https://github.com/Malfoy/PAC.

### 2.1 Prerequisites

A *set of sequences R* (also called *dataset*) is a set of finite strings on the alphabet Σ = {*A, C, G, T* }. In practice, sets of sequences can be read sets or genome sets. An input *database D* = {*R*_1_, … *R*_*n*_} is a set of *datasets*. Similarly to the datasets, a *query sequence* is a finite string from which all distinct *k*-mers are extracted (typically, a gene, a transcript, or some genomic context around a base mutation).

In the following, we suppose that broadly-used concepts in the context of computational genomics are known (namely, *k*-mers, Bloom filters [8], and minimizers [29, 12]). However, their definitions are recalled in the Appendix if needed.

#### Problem statement

The structures described hereafter estimate the cardinality of the intersection of dataset’s *k*-mers with the query’s *k*-mers. Then, according to a threshold parameter *τ*, the query is said to be *in* a dataset if its intersection is larger or equal to *τ*.

#### A sequence Bloom tree (SBT)

is an approximate membership query structure introduced in [31]. SBTs use a set of *n* Bloom filters, each representing the distinct *k*-mers of the *n* datasets in an input database. A SBT is a balanced binary tree that represents separately each Bloom filter in its leaves and the union of all Bloom filters in its root. Therefore, it allows fast membership queries in the whole database using recursive queries along the tree. We call a *sequence Bloom matrix* the matrix of Bloom filters introduced in BIGSI [10]. For SBTs, a Bloom filter is built for each of the *n* datasets in an input database. Then these filters are stacked to become a matrix in which each Bloom filter is a column. By accessing a row, one can directly know whether an element is present or absent in all Bloom filters. A *SeqOthello* is an approximate membership query structure that relies on a different paradigm. It operates as a hash table where pairs of (*k*-mer, presence/absence bitvector) are associated using a static hashing strategy and can efficiently be interrogated for *k*-mer’s presence in bit-vectors. In the following section we will define the necessary concepts to detail our novel structure.

### 2.2 Partitioned Aggregated Bloom Comb Trees construction

#### 2.2.2 Aggregated Bloom filters

PAC, as most AMQ structures, relies on Bloom filters to represent *k*-mers presence. For fixed parameters (*k, b, h*) (respectively, *k*-mer size, Bloom filter size, and number of hash functions), for each *R*_*i*_ (0 *< i* ≤ *n*) in *D*, we build a Bloom filter *BF*_*i*_ and populate *BF*_*i*_ with *R*_*i*_’s *k*-mers. Thus, each Bloom filter represents the *k*-mers of a dataset from *D*. As SBT, we organize our Bloom filters in a tree whose inner nodes result from a merge operation on Bloom filters:

##### Definition 1

(***Bloom filter merge****) Let two Bloom filters BF*_1_ *and BF*_2_ *with the same set of parameters* (*b, h*). *We define the bf merge operation as a bitwise OR:*

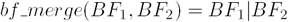

##### Definition 2

(***union Bloom filter****) A union Bloom filter is the Bloom filter result of a bf merge operation, with conserved parameters* (*b, h*).

The tree is then built by *merging* filters from bottom to top. The tree root represents the union Bloom filters of all distinct *k*-mers in *D*. PAC involves a novel tree topology in comparison to other SBTs that are balanced binary trees, which will be detailed in the following.

##### Definition 3

(***Aggregated Bloom filter****) We define a series of Bloom filters BF*_1_, *BF*_2_, … *BF*_*t*_ *built using common* (*b, h*) *parameters. They represent k-mer sets R*_1_, *R*_2_, …, *R*_*t*_ *such that R*_*t*_ ⊆ … *R*_2_ ⊆ *R*_1_. *We encode this particular series of t filters of size b as a matrix M of size t* × *b. Let V be an integer array of size b. For* 1 ≤ *i* ≤ *t, V* [*i*] *is the length of the run of 1 in the row i of M. V can effectively encode such a series of* 2^S^ *Bloom filters of size b using b integers of size S. In this way, we can encode t of such Bloom filters of size b using b* × log *t bits. We call such an integer array V endoding for the serie of Bloom filters BF*_1_, *BF*_2_, … *BF*_*t*_ *an Aggregated Bloom filter*.

##### Observation 1

*One can notice that any branch of a SBT could be represented by an Aggregated Bloom filters, with BF*_*t*_ *being the leaf node in the branch, and BF*_1_ *the root node*.

#### 2.2.2 Aggregated Bloom Comb Trees

##### Definition 4

(***Comb tree****) We call a comb tree a binary tree whose each internal node has at least one leaf as a child. When considering an order on the leaves, a* left-comb tree *(respectively*, a right-comb tree*) has its root connected to the rightmost leaf (respectively the leftmost leaf)*.

PAC relies on Bloom comb-trees to organize the set of Bloom filters representing each dataset.

##### Definition 5

(***Bloom comb tree****) We call a Bloom comb tree a comb tree built using the bf merge operation. First, given a list of Bloom filters BF*_1_, *BF*_2_, … *BF*_*n*_ *representing datasets, the leaves of the comb tree are built. Each leaf contains a Bloom filter as in SBTs. A Bloom left-comb tree is such that the leftmost inner node BF*_*n*+1_ = *bf merge*(*BF*_1_, *BF*_2_). *The right-comb tree can be defined by symmetry*.

##### Property 1

*In a Bloom left-comb tree built from a list of Bloom filters BF*_1_, *BF*_2_, … *BF*_*n*_, *we have BF*_*n*+1_ ⊆ *BF*_*n*+2_ ⊆ … *BF*_*n*+*height*_ *(with height being the height of the comb) due to the union operation performed in bf merge*.

See Figure 1 for an example of the difference between a tree used by SBTs and a Bloom comb tree.

**Figure 1:**
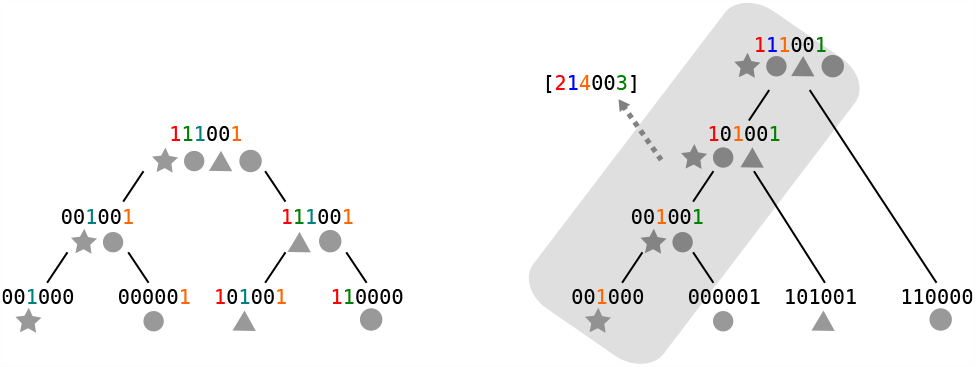
Left: a SBT structure. Right: a Bloom left comb-tree for the same database. Four datasets of a database are represented using grey shapes. Toy example Bloom filters are represented as bit vectors associated with shapes, and the content of each node is recalled. In the case of the Bloom comb tree, the inner nodes’ Bloom filters (highlighted zone in grey) are not explicitly represented but are encoded using an Aggregated Bloom filter instead (integer vector). Runs of 1’s corresponding to the integers are colored. For instance, the leftmost 1 is found at levels 1 and 2, therefore a run of length 2 in the vector.

##### Observation 2

*Similarly to the previous observation, one can notice that the longest branch of the comb is a series of Aggregated Bloom filters*.

Following Observation 2, we propose using an Aggregated Bloom filter to encode the longest branch of the Bloom comb tree.

##### Definition 6

(***Aggregated Bloom Comb Tree****) Unless otherwise specified, we denote by Aggregated Bloom Comb Tree a structure composed of a pair of Bloom left and right comb trees built on the same list of leaves. They are represented using two components. First, two Aggregated Bloom filters V*_*l*_, *V*_*r*_ *(see Definition 3) of size b, representing the branch going through all internal nodes down to the deepest leaf in both left and right-comb tree. Second, n Bloom filters are the leaves of both combs*.

Panel A of Figure 2 shows an example of two Bloom comb trees and the Aggregated Bloom filters that represent them. An Aggregated Bloom filter represents a run of 1’s in an inclusion series of Bloom filters. Thus, *V*_*l*_ (and symmetrically, *V*_*r*_) defines the maximal depth at which 1’s can be encountered at a given position in Bloom filters of the combs, i.e. the search space for a hit. Therefore, intersecting the two *V* ^′^*s* intervals refines the bounds for the search space at each position (see Panel B of Figure 2 for an example).

**Figure 2:**
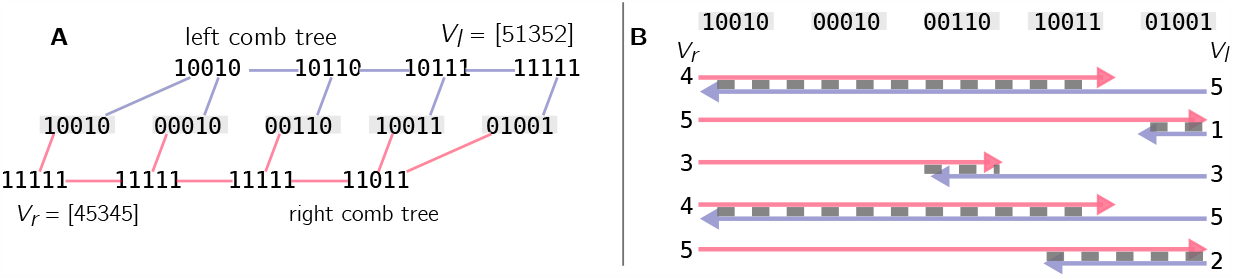
**Panel A:** Two Bloom comb trees and their Aggregated Bloom filters. The leaves Bloom filters are in the middle, colored in grey. They are the Bloom filters representing the input datasets. The left-comb tree (top) and right-comb tree (bottom) are built by aggregating the Bloom filter’s *k*-mers. The Aggregated Bloom filter *V*_*l*_ represents the leftmost path of the top comb. Respectively, *V*_*r*_ represents the rightmost path of the bottom comb. We colored in green the second bit of leaves in order to show its encoding in the Aggregated Bloom filters. In the left comb, this bit is set to 1 only in the root for the longest branch; therefore, the value is 1 (green) in *V*_*l*_. Conversely, the longest branch of the right tree has a run of 5 1’s on the second position, hence a 5 in *V*_*r*_. **Panel B:** using *V*_*l*_ and *V*_*r*_ from panel A, we show how Aggregated Bloom filters define restrained search spaces in the leaves. Again, the leaves’ Bloom filters are grey, and the values of the vectors are printed vertically. Arrows represent the intervals given by each *V*, and the final search space at each position is the dotted grey area).

#### 2.2.3 Structure partitioning

In PAC, we build a distinct *Aggregated Bloom comb tree* for each minimizer associated with a non-empty *k*-mer set.

##### Definition 7

(***super-****k****-mer [14]****) From an input string, a super-k-mer is a substring containing all consecutive k-mers that share a minimizer of size m*.

As consecutive *k*-mers in a sequence largely overlap, blocks of *k*-mers tend to share their minimizer and yield super-*k*-mers. By associating super-*k*-mers to their minimizers (and therefore the *k*-mers they come from) one can divide a *k*-mer set into up to 4^*m*^ partitions [14]. See the Supplemental Figure S1 in the Appendix for an example.

Panel A of Figure 3 shows which information is stored in a partition to represent a PAC. The combined 4^*m*^ *Aggregated Bloom comb trees* stored in 4^*m*^ partitions represent the complete *partitioned Aggregated Bloom comb tree* (PAC) structure.

**Figure 3:**
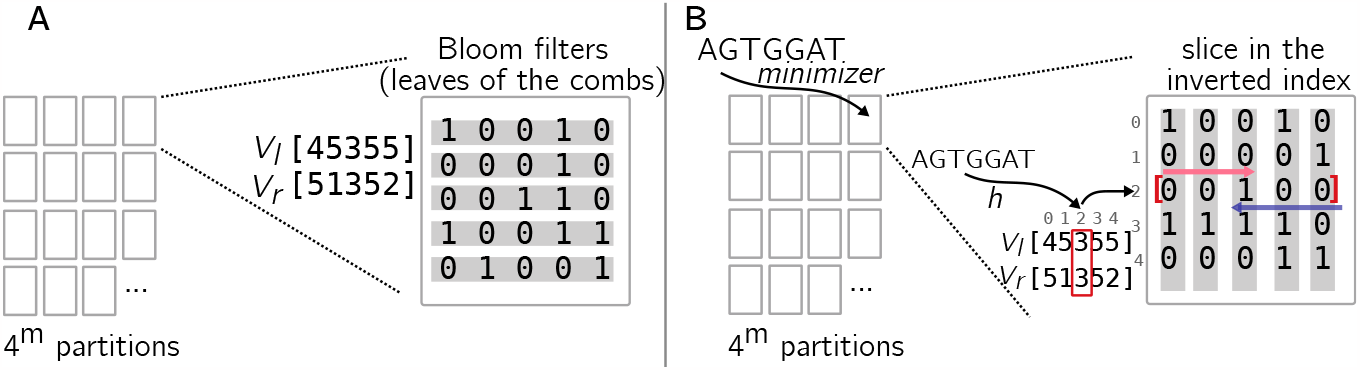
**Panel A:** the content of one partition in PAC. The Aggregated Bloom comb trees constructed for a given partition are represented using Bloom filter leaves (in gray) and two Aggregated Bloom filters *V*_*l*_ and *V*_*r*_. **Panel B:** a single query in the same partition as Panel A. The index has been inverted. A *k*-mer enters a PAC by finding the partition corresponding to its minimizer. *V*_*l*_ and *V*_*r*_ are loaded, and a search space (here [3] only) is defined given the position obtained by hashing the *k*-mer (here the *k*-mer’s hash value is 2). Then, a slice is extracted from the inverted index at the given position, and bits are checked to find 1’s. Here there’s a single position, so necessarily an 1.

#### 2.2.4 Query

##### Definition 8

(***Single query****) We call a single query the query of a single k-mer to an approximate membership query data structure. The single query ouptuts a bit-vector of size n indicating the presence / absence of a k-mer in each dataset*.

##### Definition 9

(***Multiple query****) Let s be a string of size* |*s*| *> k. We call a multiple query the query of consecutive k-mers in s through single queries to an approximate membership query data structure. The results scores are reported in an integer vector of size n*.

For all approximate membership query structures studied in this paper, the worst-case time complexity for single queries is in O(*n*). Thus, the number of random access impacts is critical for runtimes in practice.

##### Definition 10

(***Inverted Bloom filter index****) In this framework, we call an inverted Bloom filter index an index that maps each possible hash value h to a bit vector of size n that represents which filters have one at position h*.

To minimize the amount of random access, we rely on Inverted Bloom indexes during query time.

For each *k*-mer, its hash value *h* is computed, and the bit slice *h* is searched to find which Bloom filters include it. To further increase data locality, we construct one Inverted Bloom filter index per partition. In this way, querying successive *k*-mers from a super-*k*-mer is done on the same small data structure, reducing the amount of cache misses. Such partition indexes are only constructed if needed and kept to avoid redundant operations following a lazy strategy. An example is presented in Figure 3, Panel B (super-*k*-mers are omitted). Finally, we maintain and return a score vector of size *n*. It is incremented by one at a position *j* each time a *k*-mer appears in leaf *j*. Therefore, PAC is also related to Sequence Bloom matrices because queries are handled using a matrix of Bloom filters.

#### 2.2.5 Update PAC by inserting new datasets

Adding new datasets does not require changing the index structure. New datasets can be added by constructing their Bloom filters and updating the values of the Aggregated Bloom filters when needed. We implemented this functionality to allow the user to insert a dataset collection into an existing index. This insertion presents a very similar cost to building an index from the said collection, as both the construction and update algorithms consist of successive insertions. It avoids the need to rebuild the index from scratch due to novel datasets.

#### 2.2.6 Implementation details

##### Memory usage and parallelization

Bloom filters are constructed in RAM and serialized on disk to avoid heavy memory usage. In this way, only one Bloom filter is stored in RAM at a given time (*C* if *C* threads are used to treat the datasets in parallel). A PAC index is distributed into *P* sections serialized in *P* different files. Such sections can be handled separately, being mutually exclusive. This strategy provides inherent coarse-grained parallelism, as different threads can operate on different sections without mutual exclusion mechanisms during construction. It also grants low memory usage as only small sections (i.e., a fraction of the total index) have to be stored in memory at a given time.

Similarly, the query also benefits from partitioning. All *k*-mer associated with a given partition are queried at once in a sequential way to limit memory usage. Each Bloom filter is loaded separately in RAM and freed after the inverted index is constructed to be queried (*C* Bloom filters can be loaded simultaneously in RAM if *C* threads are used). This behavior guarantees that each partition will only be read from the disk once and that only one partition will be stored in RAM at a given time. Furthermore, partitions not associated with any query *k*-mer can be skipped. Both construction and query benefit from improved cache coherence as several successive *k*-mers are located in the same small structure, generating cache misses for groups of *k*-mers instead of doing so for nearly each *k*-mer.

##### Inverted index

Bloom filters are represented as sparse bit-vectors (using the BitMagic library^1^) in order to optimize memory usage.

## 3 Results

All experiments were performed on a single cluster node running with Intel(R) Xeon(R) Gold 6130 CPU @ 2.10GHz with 128GB of RAM and Ubuntu 22.04.

### 3.1 Indexing 2,500 human datasets and comparison to other approximate membership query structures

We compared PAC against the latest methods of the approximate membership query paradigm, i.e., HowDeSBT [16] for SBTs and SeqOthello [33]. We also tested COBS [6], the most recent sequence Bloom matrix, although it is not designed to work with sequencing data, but rather genomes. We used kmtricks [20] for an optimized construction of HowDeSBT and its Bloom filters (commit number 532d545). SeqOthello (commit 68d47e0) uses Jellyfish [22] for its pre-processing, we worked with version 2.3.0. COBS’s version was v0.1.2, and PAC’s commit was cee1b5c. We used the classic mode of COBS that does not rely on folding to ensure a fair comparison of Bloom filter size and pratical false positive rate accros tools.

The dataset first used in the initial SBT contribution [31] has become a *de facto* benchmark in almost any subsequent article that describes a related method. We make no exception, and bench-marked PAC and other approximate membership query structures on this dataset. It contains 2,585 human RNA-seq datasets, with low-frequency *k*-mers filtered according to previous works [32]^2^, resulting in a total of ∼ 3.8 billion of distinct *k*-mers. The input files are represented as compacted de Bruijn graphs generated by Bcalm2 [13] in gzipped FASTA files. We also kept the settings used in previous benchmarks and used a *k* value of 21. We chose COBS, PAC, and HowDeSBT’s settings to have an average 0.5% false positive rate. Notably, SeqOthello’s false positive rate cannot be controlled.

In Table 1, we present the costs of indexing this database with the different tools. Several observations can be made. First, PAC does not generate any temporary disk footprint; only the final index is written on disk, while HowDeSBT, SeqOthello, and COBS generate many temporary files that are an order of magnitude larger than the index itself. Another observation is that PAC and COBS’s pre-processing and index construction steps are comparable and faster than the other tools, showing at least a two-fold improvement of required CPU time. The total computational cost of a PAC index is improved three times over SeqOthello and six times over HowDeSBT. Memory-wise, the memory footprint of HowDeSBT and PAC are both low, while SeqOthello and COBS are somewhat higher without being prohibitively high. Finally, excluding COBS, the different produced indexes are of the same order of magnitude, even if PAC presents the heaviest index, slightly larger than SeqOthello, while HowDeSBT is the smallest. This can be explained by the fact that tree based tools (as HowDeSBT) spend a lot of time to chose how to group and merge files to optimize the index compressibility resulting in a hard to construct but smaller indexes tradeoff. Comptuting a efficient file ordering to boost compression is an interesting problem that would benefit any matrix based index. Being compression-free, COBS’ index is two orders of magnitude larger than the other tools and produces even larger temporary files. COBS’ CPU time is similar to PAC’s, but its heavy disk usage presents a higher wall-clock time on our hard-disk drives.

**Table 1:** Index construction resource requirements on 2,585 RNA-seq. The column “Max temporary disk” reports the maximal external space taken by the method (usually during preprocessing). The “Final Index Size” indicates the final index size when stored on disk. The execution times of the methods are reported in CPU hours. All methods were run with 12 threads.

| Tool | Max temporary disk (GB) | Preprocessing time (CPU,h) | Index construction time (CPU,h) | Peak RAM (GB) | Final index Size (GB) |
| --- | --- | --- | --- | --- | --- |
| COBS | 4,914 | **7** | **1** | 111 | 2,458 |
| HowDeSBT | 650 | 16 | 44 | **4.2** | **15** |
| SeqOthello | 1,000 | 19 | 12 | 43 | 25 |
| **PAC** | **0** | 8 | **1** | 9.6 | 28 |

### 3.2 Indexing over 32,000 human datasets and scalability regarding the number of datasets

We downloaded 32,768 RNA-seq samples from SRA (accession list in available on the github repository^3^) for a total of 30 TB of uncompressed FASTQ data. We created incremental batches of 2^8^, 2^9^, …, up to 2^15^ datasets to document the scalability of methods according to the number of datasets. In this second experiment, the indexes are built directly on sequencing datasets in raw FASTA format, containing redundant *k*-mers and sequencing errors. Each tool was configured to filter unique *k*-mers before indexing to remove most sequencing errors (we used *k* = 31). Since COBS does not provide such an option and produces huge index files, we did not include it in this benchmark. To approximate the order of magnitude of the amount of (non-unique) distinct *k*-mers, we used ntcard [25]. For example, the batch of 2^11^ datasets contains approximately 3 billion *k*-mers to index, while the 2^14^ dataset contains more than 40 billion *k*-mers.

In Figure 4, we report the CPU time required for index construction (including preprocessing) to display their evolution on databases of increasing size. In Figure 5, we report temporary disk usage during index construction and the final index sizes on the disk after compression.

**Figure 4:**
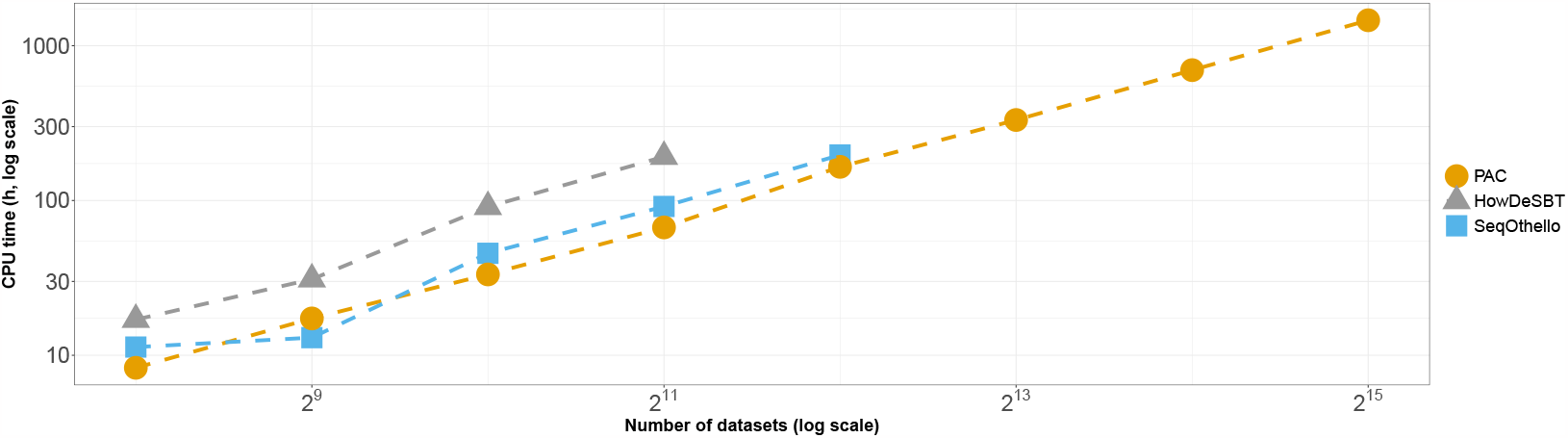
Results on increasing human dataset sizes. We report total CPU hours for constructing the different indexes, including the Bloom filter construction and index constructions. X-axis is in log 2 scale and Y-axis is in log 10 scale. All methods were run with 12 threads.

**Figure 5:**
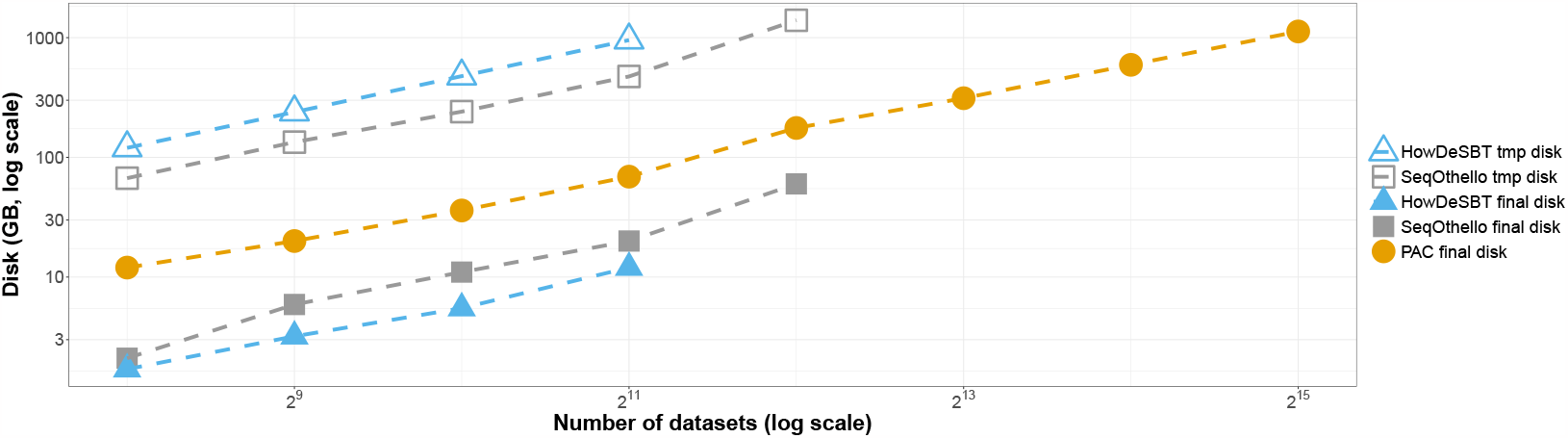
Results on increasing human dataset sizes. We report disk footprints, both temporary and final, for constructing the different indexes. Note that PAC does not use temporary disk. X-axis is in log 2 scale and Y-axis is in log 10 scale. All methods were run with 12 threads.

We can make observations similar to those in the first experiments. PAC is the only tool that managed to build an index of the 32,000 human RNA-seq samples, in ∼5 days (∼60 CPU days), using 22GB of RAM and for a total index size of 1.1TB. PAC’s construction is faster than the state-of-the-art for the whole construction process. It produces slightly bigger indexes, but requires much less external memory. SeqOthello crashed during index construction in the experiment with 2^12^ datasets, and HowDeSBT failed to end before our ten-day timeout in the experiment with 2^11^ datasets. PAC was the only tool to build an index on the 2^13^, 2^14^, and 2^15^ datasets.

### 3.3 Indexing microbial datasets

Besides indexing RNA-seq data, approximate membership query indexes were also used to index large collections of bacterial genomes. To further highlight the scalability of PAC, we built it on two massive bacterial collections. In a first experiment, we fetched the 661,000 bacterial genomes representing more than 3 Tera bases, which were previously collected in a database [7] and built two indexes on this collection. A first index featuring large Bloom filters (2^29^ bits) for low *k*-mer query false-positive rates (below 1% for 5 Mega-bases genomes) and a second index with smaller Bloom filters (2^27^ bits) when higher false-positive rates can be allowed (below 7% for 5 Mega-bases genomes). Both indexes were constructed within 24 hours using 21 and 6 GB of RAM, respectively, for a total size of 2.4 and 1.4 TB, respectively.

In a second experiment, we downloaded all bacterial assemblies available on Genbank (counting 1,200,575 genomes at the time of the experiment and representing more than 5 Tera-bases) and built an index with a Bloom filter size of 2^27^ bits that corresponds to a false positive rate around 4% for 5 Mega-bases genomes. The construction lasted 24 hours for a total of 410 CPU hours using less than 5GB of RAM for a total index size of 3.5 TB. This is to our knowledge the largest collection ever indexed by an approximate membership query.

### 3.4 Query results on human RefSeq

We designed a similar experiment to the one presented in HowDeSBT’s paper, with sequence batches of increasing sizes. We repetitively selected random transcripts from human RefSeq (using seqkit [30]), and gathered 10, 10, 5, 3 and 3 batches of sizes 1, 10, 100, 1,000 and 10,000 transcripts. We report HowDeSBT, SeqOthello, and PAC’s CPU time on each batch size in Figure 6. We warmed the cache before profiling the queries. In accordance with the literature [16], we observe that Sequence Bloom Tree structures perform the best on small query sets. In larger instances, other methods can be preferred. SeqOthello shows the best performance and keeps a relatively constant query time over the input size. PAC has the same behavior, while being slightly more CPU time than SeqOthello. We also note that no method required more than 10GB of RAM to perform the queries. PAC had the lowest RAM footprint, with up to 2GB for 10,000 transcripts. COBS CPU time usages are not representative of its actual queries’ performance, as it incurs almost no CPU times (typically less than 1 CPU second per transcript), so we chose not to plot it along with its competitors.

**Figure 6:**
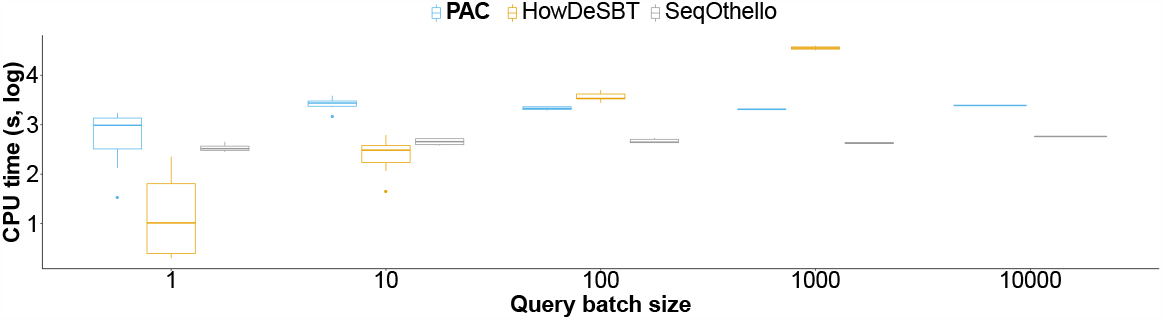
Results on query batches. We present query CPU times computed on batches of 1 to 10k transcript sequences. The Y-axis is on a log 10 scale. The queries were computed using 12 threads.

We performed a separate benchmark to assess the queries wall-clock times. On a batch of 100 transcripts, HowDeSBT lasted 1h30, COBS lasted 30 minutes, SeqOthello lasted 10 minutes, and PAC lasted 5 minutes. Interestingly, on a larger batch of 10,000 transcripts, SeqOthello and PAC presented very similar results (10 and 5 minutes, respectively). These results indicate that the query time of these two tools is dominated by index loading. To assess when query time became predominant, we queried a large batch of 100,000 transcripts for which PAC lasted 11 minutes (2 CPU hours) and an even larger batch of 500,000 transcripts handled by PAC in 40 minutes (6.5 CPU hours).

### 3.5 Impact of comb structure on query for collections entailing high *k*-mer diversity

One novelty of PAC is the use of lightweight Aggregated Bloom filters to accelerate the queries by skipping subsections of the bitslices. To assess the efficiency of this strategy, we report in Table 2 the query times of several query batches on a index composed of all complete *S. enterica* genomes from RefSeq (counting 11,993 genomes at the time of the experiment). We report three different query times for each batch, using, respectively, two Aggregated Bloom Comb trees (according to definition 6), a single one and none (i.e., the structure is a Bloom matrix). Without the use of Aggregated Bloom filters PAC index is conceptually identical to a matrix approach such as COBS while using one or two Aggregated Bloom filters allow to skip some bitslice of the index. The number of bits that can be skipped depends in practice on the similarity shared between the query and the indexed documents. To highlight this effect, a batch is made up of *S. enterica* that are highly similar to the indexed genomes, a batch is made up of *E. coli* that are very dissimilar to the indexed genomes, and a batch is made up of random sequence to show an extreme example of dissimilarity. The first observation is that Aggregated Bloom filters hardly improve the query time of the *S. enterica* batch. This result is expected because most query *k*-mers should be found in many indexed genomes and the amount of zeros that could be skipped in such slices should be very low. However, we see that using one Aggregated Bloom filter greatly accelerates the query on the two dissimilar batches, increasing the throughput by several fold. Furthermore, using a second Aggregated Bloom filter accelerates the query time even more but is not as beneficial as the first one.

**Table 2:** Results on query times in CPU hours according to the PAC mode used along with the speedup obtained compared to the classical matrix approach (No ABF column). Double, Single, No ABF denotes the number of combs used in the structure. We present query CPU times computed along with the overhead constant across modes that include parsing the query file and loading the index. The random genomes are random nucleotide sequences of length 5 megabases. *E*.*coli* and *S*.*enterica* genomes were randomly selected among the RefSeq genomes.

| Query Dataset | Double ABF filter | Single ABF filter | No ABF | Overhead |
| --- | --- | --- | --- | --- |
| 200 Random genomes | 0.7 (13.6x) | 1.3 (7.3x) | 9.5 | 1.0 |
| 200 *E.coli* genomes | 1.2 (7x) | 2.2 (3.8x) | 8.4 | 1.0 |
| 200 *S.enterica* genomes | 10.4 (1.2x) | 11.6 (1x) | 12 | 1.0 |

## 4 Discussion

To our knowledge, PAC is the first approximate membership query *k*-mer set structure to index the entire Genbank bacterial genome collection and reach 32k human RNA-seq datasets.

PAC combines the simplicity and efficiency of inverted index matrix approaches such as COBS or BIGSI with a lightweight tree structure.

The novel tree structure has a minimal resource footprint, yet greatly improves the query time when a query is dissimilar to the index content, a scenario possibly met with microbial databases. As *k*-mer sets are designed to efficiently skip unrelevant documents, our Aggregated Bloom filters allow us to efficiently prune our query space.

Using several real datasets, we demonstrate that PAC is practically scalable for its construction. In contrast to other approaches, PAC is simultaneously frugal in RAM, disk, and time requirements for building an index.

We showed PAC’s ability to query 500,000 human transcripts in less than an hour, being the fastest in wallclock time, and comparable to SeqOthello in CPU time. The worst-case query complexity remains the same for all methods, O(*n*) for a *k*-mer present in all *n* datasets.

We reviewed that inverted indexes methods perform Θ(1) random accesses, but still need to read bit-slices of size O(*n*). Using Aggregated Bloom filters, PAC improves these indexes, as in favorable cases (*k*-mers present in a single dataset with no collision or absent everywhere), PAC can answer in O(1).

## 5 Competing interests

No competing interest is declared.

## 6 Acknowledgments

The authors would like to thank Léonid and Zora for a renewed sense of what tiredness, teamwork and joy are. We also thank Daniel Gautheret for providing us with tips on the SRA API, RECOMB-Seq reviewers, and Anatoliy Kuznetsov for his outstanding Bitmagic library and sustained support. This work was supported by grants from the Agence Nationale de la recherche for the project “full-RNA “ [ANR-22-CE45-0007)] and”AGATE “ [ANR-21-CE45-0012].

## 7 Appendix

### 7.1 Definitions

A *Bloom filter* (BF) is a space-efficient data structure for inexact set representation. For Bloom filters, inexact means that queries to such a representation have a controlled rate of false positives and no false negatives. A Bloom filter has two parameters (*b, h*), which define the size of a bitvector and the number of hash functions of the filter. A Bloom filter maps each set element in the bitvector using *h* hash functions, and sets mapped bits to 1 (initially, all bits are 0’s). The query follows the same principle and verifies that all hashed locations contain a 1. False positives on foreign keys occur because of hash collisions (which can be controlled by tuning (*b, h*)).

A *minimizer [29]* of size *m* of a *k*-mer *S* (*m < k*) is the smallest hashed substring of size *m* in *S*.

### Proof of observation 1

We define a series of Bloom filters *BF*_1_, *BF*_2_, … *BF*_*t*_ built using common (*b, h*) parameters. They represents *k*-mer sets *R*_1_, *R*_2_, …, *R*_*t*_ such that *R*_*t*_ ⊆ … *R*_2_ ⊆ *R*_1_. We encode this particular series by viewing it as a matrix *M* of size *t* × *b*. We recall that a SBT is built using the merge operation on pairs of leaves to obtain a binary tree.

**Proof 1** *One can notice that by recursion, any bit set to 1 in BF*_*i*_, (1 *< i* ≤ 2) *is also set to 1 in BF*_*i*-1_, … *BF*_1_. *Similarly, if a bit is set to 1 in a Bloom filter of a SBT, the 1 is conserved through the recursive OR operations that build the parent nodes of this Bloom filter*.

#### 7.1.1 Implementation details

##### *K*-mer representation

Each *k*-mer is represented by the lexicographically smallest of the forward string and reverse-complemented string (canonical *k*-mer) and is hashed using a fast xorshift hashing function.

Figure S1 presents the partition scheme based on minimizers and super-*k*-mers.

**Figure S1:**
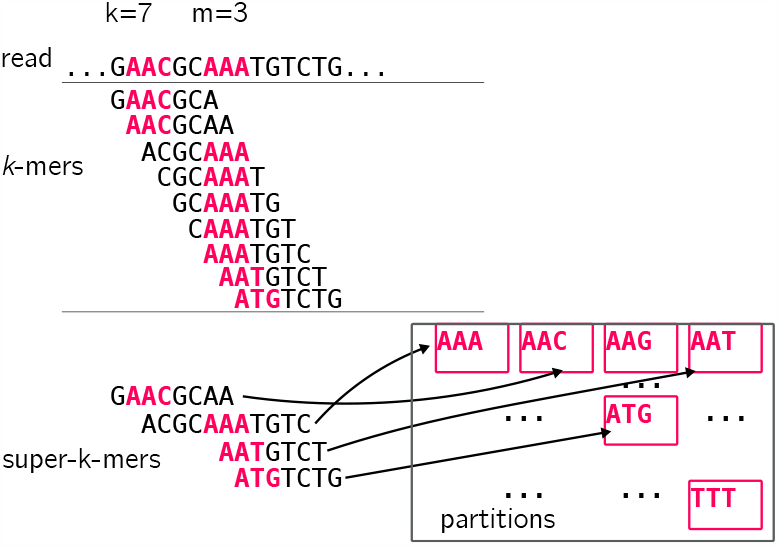
Example of minimizers and super-*k*-mers computed from a read (*k* = 7, *m* = 3). Minimizers are represented in pink (for the sake of simplicity, we consider the lexicographic order on minimizers). See how the second super-*k*-mer aggregates several *k*-mers having AAA as a minimizer. We show partitions corresponding to minimizers in the same color.

## Footnotes

1 https://github.com/tlk00/BitMagic

2 https://www.cs.cmu.edu/~ckingsf/software/bloomtree/srr-list.txt

3 https://github.com/Malfoy/PAC/blob/main/listpacpaper.txt.gz

## Notes

### Competing Interest Statement

The authors have declared no competing interest.

### Summary of Updates

new results

